# Delamanid and pretomanid have comparable bactericidal activity but pretomanid potently inhibits *Mycobacterium tuberculosis* ribosomal rRNA synthesis

**DOI:** 10.64898/2025.12.30.697025

**Authors:** Matthew J. Reichlen, Emmanuel Musisi, Samuel T. Tabor, Holly Nielsen, Ashley M. Gerwing, Firat Kaya, Matthew Zimmerman, Martin I. Voskuil, Gregory T. Robertson, Nicholas D. Walter

## Abstract

**Background:** The nitroimidazoles delamanid and pretomanid play an important role in contemporary tuberculosis treatment. It is unclear whether delamanid and pretomanid have meaningfully different activity since both reduce *Mycobacterium tuberculosis* colony forming units (CFU) similarly in animal models. The RS ratio is a pharmacodynamic marker of ongoing rRNA synthesis that has been associated with treatment-shortening (*i*.*e*., sterilizing) activity.

**Methods:** Using *Mycobacterium tuberculosis Erdman*, we conducted dose-ranging studies in aerobic axenic culture and in the conventional BALB/c mouse high-dose aerosol infection model to compare bactericidal and RS ratio activity of delamanid and pretomanid.

**Results:** *In vitro* concentration-response curves showed that delamanid and pretomanid had similar RS ratio effect at maximal concentration but pretomanid was more potent, achieving 90% of the maximal effect (RS-EC_90_) at a lower concentration (390 ng/mL) than delamanid (810 ng/mL). In mice, delamanid and pretomanid had similar effects on CFU. Human-equivalent doses of delamanid (6 mg/kg) and pretomanid (50 mg/kg) resulted in plasma C_max_ concentrations well below (210 ng/mL) and well above (7,825 ng/mL) the RS-EC_90_, respectively. Delamanid displayed no discernable RS ratio response, even at 16-times the human-equivalent dose. Higher pretomanid doses resulted in significantly greater RS ratio effects.

**Conclusions:** We found that delamanid and pretomanid have similar bactericidal activity but pretomanid has superior RS ratio activity. Meaningful differences between drugs within the same class were not captured by conventional CFU-based pharmacodynamics, supporting the value of measuring orthogonal drug effects such as the RS ratio.

**LAY SUMMARY:** Antibiotics in the nitroimidazole class are used in treatment of drug-resistant tuberculosis. There are two approved nitroimidazole antibiotics: delamanid and pretomanid. For decades, it has been unclear whether delamanid and pretomanid are interchangeable or whether they affect the bacterium *M. tuberculosis* differently. Most studies of the effect of antibiotics count the number of bacterial colonies that form on a culture plate. “Colony forming units” tell us about change in bacterial burden but does not give information about bacterial health. A new way of thinking about antibiotic effect is the RS ratio. The RS ratio is a test that measures how much ribosomal RNA synthesis is ongoing. Ribosomal RNA synthesis is a “vital sign” of bacterial health and activity. The key finding of this study is that although the two nitroimdazole antibiotics look the same in terms of their effect on bacterial burden, they have different effects on bacterial health. This information deepens understanding of differences between two clinically important antibiotics. It also shows that antibiotics testing should consider not only bacterial burden but also new tests of bacterial health.

## INTRODUCTION

There is an urgent need for shorter, more effective tuberculosis (TB) treatments. An important new class of anti-TB drugs is the nitroimidazoles. Two nitroimidazoles, delamanid and pretomanid, received US Food and Drug Administration approval for drug-resistant TB in 2014 and 2019, respectively, and pretomanid is a component of World Health Organization recommended treatment of multiple-drug resistant TB.^1^

Despite intensive preclinical and clinical evaluation, there remains considerable uncertainty about whether delamanid and pretomanid have a meaningfully different activity or are interchangeable.^2^ As comprehensively reviewed recently, delamanid and pretomanid are structurally similar and share a dual mode of action which includes inhibition of mycolic acid synthesis and respiratory poisoning. However, delamanid inhibits two mycolic acid classes (ketomycolates and methoxymycolates) while pretomanid inhibits only one (ketomycolates).^3–5^ Additionally, while both appear to poison *M. tuberculosis* (*Mtb*) respiration, delamanid does so via an NAD-adduct^6^ while pretomanid generates reactive nitrogen.^7,8^ When administered at an identical dose in murine models, delamanid exhibits greater bactericidal activity (*i*.*e*., greater reduction in colony forming units (CFU)) than does pretomanid. However, when given at lower doses designed to mimic human AUC, the bactericidal activity of delamanid and pretomanid are similar.^2^ Because pre-clinical studies have not clearly differentiated the activity of delamanid and pretomanid, there has been persistent uncertainty regarding optimal usage in humans. As a result, considerable resources are currently being expended to support head-to-head testing of delamanid versus pretomanid in human clinical trials.^9,10^

One reason the activity of delamanid and pretomanid have been difficult to differentiate in preclinical studies is that efficacy has generally been estimated in terms of CFU. CFU enumerates the burden of *Mtb* capable of growth on agar but provides little information about the effect of drugs on *Mtb* physiology. In the conventional BALB/c high-dose aerosol (HDA) mouse infection model, which is the workhorse of *in vivo* efficacy testing, early bactericidal activity does not reliably predict time to achieve durable cure.^11^ Therefore, long-term murine relapse studies are conventionally used to quantify treatment shortening in terms of treatment time required to prevent relapse in 95% of mice (T_95_).^12^

An alternative to CFU is measuring “pathogen health,” meaning how drugs affect *Mtb* physiology.^13–18^ Drugs or regimens that reduce CFU to equivalent degrees may force unique patterns of bacterial injury and physiologic adaptation.^16,17^ One measure of pathogen health is the RS ratio^®^ assay which quantifies ongoing *Mtb* rRNA synthesis.^17^ The capacity of drugs and regimens to rapidly and profoundly suppress the RS ratio in mice has been associated with their ability to more rapidly achieve non-relapsing cure (*i*.*e*., treatment shortening).^17^ Here, we used both CFU and RS ratio to compare the effects of delamanid and pretomanid *in vitro* and in the BALB/c HDA mouse model.

## METHODS

### In vitro experiments

Cultures of *Mtb* Erdman were propagated in 7H9 medium containing 850 mg/L NaCl, 0.2% glycerol, 0.2% glucose, 0.5% BSA, and 0.05% tween80. *Mtb* cultures were grown to mid-log and then transferred into sterile glass test tubes (20 by 125 mm) containing a 12 by 4.5 mm stir bar at an OD_600_ = 0.05 and a volume of 5 mL. Cultures were agitated at ∼200 rpm under the control of a rotary magnetic tumble stirrer at 37°C in 5% CO_2_. Sterile 1000X stocks of delamanid and pretomanid prepared in DMSO were added after 18h of outgrowth. An initial 5-fold dilution series starting from 10 μg/mL of each drug was performed to determine an approximate concentration at which each drug achieved 90% of their respective RS-E_max_ values (RS-EC_90_). Along with this, a concentration 10-fold higher than the observed MIC value for each drug in this model, 16 μg/mL and 4 μg/mL for delamanid and pretomanid respectively, was also included. After which, a 2-fold dilution series starting from 8X the RC-EC_90_ was performed starting from 6.4 μg/mL and 3.2 μg/mL for delamanid and pretomanid respectively. RNA was collected from cultures both prior to drug exposure and after 48h of drug exposure as described^14^. RNA collection, isolation, and ddPCR were performed as described ^14,17^. RS ratio effect dose response was defined using an E_max_ 4-parameter variable slope sigmoid response model as described ^14^. Dose response curves were generated and analyzed using GraphPad Prism 9. Curves were fit to the observed data using least squares regression and medium convergence criteria (maximum iterations = 1000). RS ratio percent effect corresponded to the percent decrease in RS ratio after 48h drug exposure relative to controls prior to drug exposure.

### Murine experiments

Animal studies were performed at Colorado State University (CSU) in ABSL-3 containment in accordance with CSU Institutional Animal Care and Use Committee (reference number: 4179) guidelines. Six-to eight-week-old female pathogen-free BALB/c mice (Jackson Laboratories) were infected by aerosol (Glas-Col) with *Mtb* Erdman to achieve deposition of ∼3.7 log_10_ CFU in the lungs one day following high-dose aerosol infection (day -10).^19^ Treatment by oral gavage, five days per week was initiated 11 days post aerosol (day 0). Pretomanid (Chemshuttle) was prepared in the CM-2 formulation as previously described.^19^ Delamanid (ChemShuttle) was prepared in 5% gum Arabic (Sigma). Prepared pretomanid or delamanid were administered 5 of 7 days per week by oral gavage in 0.2 mL volume quaque die (QD; once per day) at 10, 50, or 100 mg/kg and 6, 10, 50, and 100 mg/kg, respectively. Mice were humanely euthanized on day 0 as a pre-treatment control, and on treatment days 5, 12 and 26. Lungs were flash frozen under liquid nitrogen for RNA and CFU enumeration. Details on RS ratio assessment were described previously.^13,17^ For CFU, left, inferior, and post-caval lung lobes (2/3rds by weight) were homogenized using the Bertin Precellys CKMix50-7 mL lysis kit [in 4.5 mL phosphate-buffered saline (PBS) with 10% bovine serum albumin (BSA)]. Lung homogenates were plated as serial dilutions on 0.4% charcoal-supplemented 7H11 agar (i.e., Middlebrook 7H11 agar plates supplemented 0.2% [v:v] glycerol, 10% [v:v] oleic acid-albumin-dextrose-catalase (OADC) supplement, and 0.01 mg/mL cycloheximide, and 0.05 mg/mL carbenicillin). For RNA, flash frozen superior and middle lung lobes were homogenized using the Bertin Precellys CKMix50-7 mL lysis kit in 1.5 mL CAMM-RPS buffer. ^13,17^

### Statistical analysis

The RS ratio effect at maximal concentration (RS-E_max_) and concentration required to achieve 90% of the maximal attainable effect (RS-EC_90_) were calculated as previously described.^20^ All *p-*values for pairwise comparisons were calculated using the Wilcoxon rank-sum test. Results with p < 0.05 were considered statistically significant. Analyses and graphics were conducted using R version 4.5.1 (R Development Core Team, Vienna, Austria https://www.r-project.org) and RStudio version 2025.05.1+513 (https://posit.co).

## RESULTS

### RS ratio activity in vitro

In *in vitro* dose-response curves,^14^ delamanid and pretomanid had indistinguishable RS ratio effects at maximal concentration (RS-E_max_), indicating equivalent RS ratio efficacy (**Fig 1a-b)**. However, the dose required to achieve 90% of the maximal attainable effect (RS-EC_90_) was higher for delamanid (810 ng/mL) than for pretomanid (390 ng/mL). Achieving the same effect at a lower concentration indicates that pretomanid has more potent RS ratio activity.

**Figure 1:**
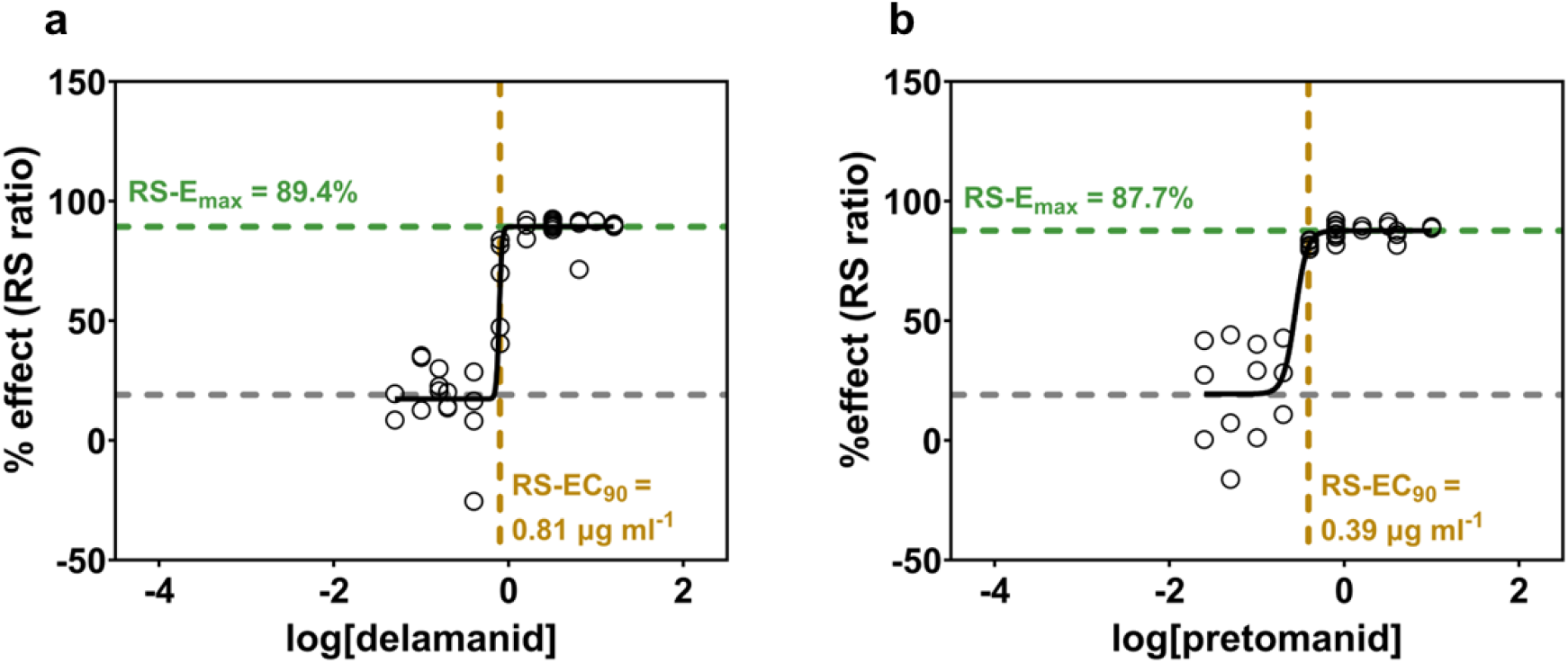
*In vitro* RS ratio dose-response sigmoid E_max_ models of delamanid and pretomanid. Solid black dashed lines depict best-fit curves. Green horizontal dashed lines depict the RS-E_max_. Grey horizontal dashed lines depict mean RS ratio response for no drug at 48 h. Yellow vertical dashed lines depict RS-EC_90_. Circles represent values from individual samples. Data for pretomanid was published previously in ref ^14^. Dose-response curves were fitted to observed data using least squares regression and medium convergence criteria (maximum iterations = 1000).

### Dose ranging in BALB/c mice

In a murine dose ranging study, no dose of delamanid reached the *in vitro* delamanid RS-EC_90_ of 810 ng/mL (**Fig 1a, Fig 2a**). For example, the 100 mg/kg delamanid dose (*i*.*e*., >16-times higher than the established human equivalent 6 mg/kg dose) resulted in a C_max_ of 707 ng/mL.

**Figure 2.**
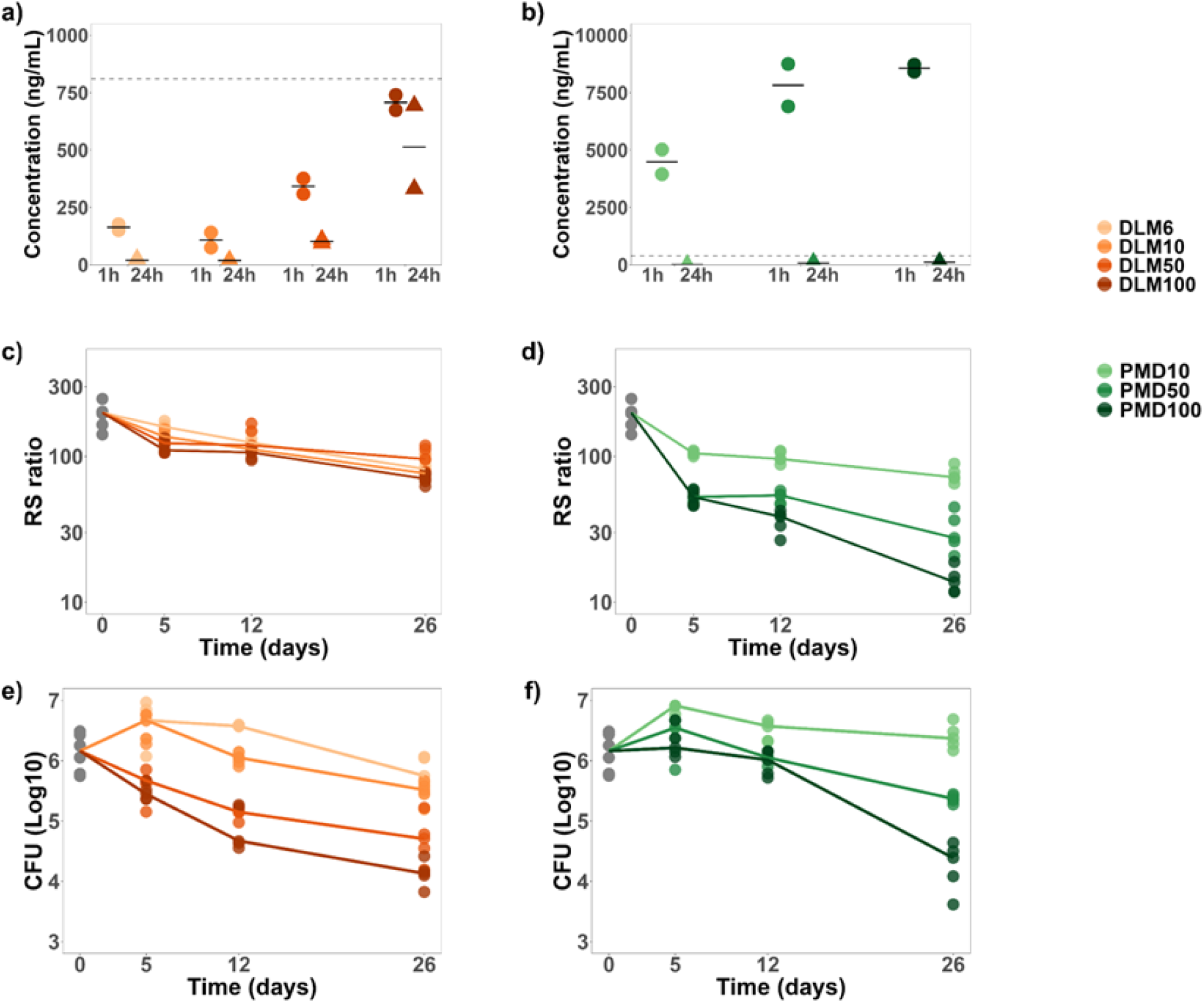
Pharmacokinetics and pharmacodynamic markers during treatment with varying concentrations of delamanid and pretomanid (**a–b**) C_max_ and trough concentrations following treatment with varying doses of (**a**) delamanid and **(b**) pretomanid after 1 h (circles) and 24 h (triangles). Horizontal bars indicate median values. Dotted gray lines indicate the RS-EC_90_ concentration determined *in vitro*. (**c–d)** Change in RS ratio during treatment with varying doses of (**c**) delamanid and (**d**) pretomanid. (**e–f**) Change in CFU during treatment with varying doses of (**e**) delamanid and (**f**) pretomanid. For **c-f**, circles represent values from individual mice and lines connect median values.

By contrast, the standard human-equivalent pretomanid dose (50 mg/kg) resulted in a C_max_ of 7,825 ng/mL, far exceeding the *in vitro* pretomanid RS-EC_90_ of 310 ng/mL (**Fig 1b, Fig 2b**). At the standard human equivalent dose of 50 mg/kg, the trough pretomanid concentration was 78 ng/mL. The highest dose of pretomanid (100 mg/kg) resulted in higher C_max_ and trough concentrations than standard dosing although the trough concentration remained lower than the *in vitro* pretomanid RS-EC_90_.

### RS ratio activity in mice

Consistent with the lower-than-RS-EC_90_ plasma concentrations described above, delamanid had no discernable RS ratio dose-response relationship in mice (**Fig 2c**). By contrast, increasing pretomanid doses resulted in progressively greater decreases in the RS ratio (**Fig 2d**). When compared at human-equivalent doses (*i*.*e*., 6 mg/kg and 50 mg/kg for delamanid and pretomanid, respectively), pretomanid had significantly greater RS ratio activity than delamanid (*P=*0.008 at all timepoints). When delamanid and pretomanid were given at an identical dose (resulting in a delamanid dose substantially exceeding human exposures), pretomanid continued to have significantly greater RS ratio activity than delamanid (**Table 1**). For example, when both drugs were given at a dose of 50 mg/kg, pretomanid had significantly greater RS ratio activity than delamanid (*P*=0.008 at all treatment days).

**Table 1.** Median RS ratio at various durations and doses of delamanid and pretomanid. Wilcoxon *P-*values comparing equal dosing are shown.

| Day/Dose | Median RS ratio |  | *P*-value |
| --- | --- | --- | --- |
|  | Delamanid | Pretomanid |  |
| Day 5 |  |  |  |
| 6 mg/kg<sup>*</sup> | 159.3 | -- | --- |
| 10 mg/kg | 136.4 | 104.9 | 0.008 |
| 50 mg/kg<sup>¶</sup> | 123.0 | 52.7 | 0.008 |
| 100 mg/kg | 110.2 | 52.5 | 0.008 |
| Day 12 |  |  |  |
| 6 mg/kg<sup>*</sup> | 124.2 | -- | --- |
| 10 mg/kg | 111.6 | 95.9 | 0.055 |
| 50 mg/kg<sup>¶</sup> | 118.9 | 54.0 | 0.008 |
| 100 mg/kg | 106.3 | 38.6 | 0.008 |
| Day 26 |  |  |  |
| 6 mg/kg<sup>*</sup> | 81.9 | -- | --- |
| 10 mg/kg | 76.4 | 95.9 | 0.31 |
| 50 mg/kg<sup>¶</sup> | 95.7 | 38.6 | 0.008 |
| 100 mg/kg | 70.2 | 38.6 | 0.008 |

<sup>\*</sup> Standard human-equivalent doses [plasma AUC] are 6 mg/kg delamanid, and <sup>¶</sup> 50 mg/kg for pretomanid.

### Bactericidal activity in mice

Both delamanid and pretomanid displayed a dose-response relationship in their effect on CFU (**Fig 2e-f**). Consistent with previous studies, delamanid and pretomanid had indistinguishable bactericidal activity when compared at respective human-equivalent doses at day 5 (*P*=0.75) and day 26 (*P*=0.15). At day 12, the human-equivalent pretomanid dose decreased CFU marginally but significantly more than the human-equivalent delamanid dose (*P*=0.04). When delamanid and pretomanid were given at the same dose, delamanid generally had significantly greater bactericidal activity (**Table 2**). For example, when both drugs were given at 50 mg/kg, delamanid reduced CFU significantly more than pretomanid at day 12 (*P=*0.01) and at day 26 (*P*=0.008).

**Table 2.** Change in CFU relative to pre-treatment control (log_10_). Positive and negative values indicate an increase or decrease in CFU relative to pre-treatment control, respectively. Wilcoxon *P-*values comparing equal dosing are shown.

|  | Decrease in CFU (log<sub>10</sub>) |  |  |
| --- | --- | --- | --- |
| Day/Dose | Delamanid | Pretomanid | *P*-value |
| Day 5 |  |  |  |
| 6 mg/kg<sup>*</sup> | + 0.51 | --- |  |
| 10 mg/kg | + 0.51 | + 0.75 | 0.18 |
| 50 mg/kg<sup>†</sup> | - 0.49 | + 0.38 | 0.065 |
| 100 mg/kg | - 0.71 | + 0.05 | 0.002 |
| Day 12 |  |  |  |
| 6 mg/kg<sup>*</sup> | + 0.41 | --- |  |
| 10 mg/kg | + 0.11 | + 0.41 | 0.03 |
| 50 mg/kg<sup>†</sup> | - 1.01 | - 0.11 | 0.002 |
| 100 mg/kg | - 1.49 | - 0.11 | 0.009 |
| Day 26 |  |  |  |
| 6 mg/kg<sup>*</sup> | - 0.41 | --- |  |
| 10 mg/kg | - 0.65 | + 0.21 | 0.004 |
| 50 mg/kg<sup>†</sup> | - 1.46 | - 0.79 | 0.04 |
| 100 mg/kg | - 2.03 | - 1.78 | 0.59 |

\* Standard human-equivalent doses are 6 mg/kg for delamanid, and <sup>†</sup> 50 mg/kg for pretomanid.

## DISCUSION

Our *in vitro* and murine studies revealed a hitherto unreported difference between the effects of delamanid and pretomanid. Although the two drugs had similar bactericidal activity, we found that pretomanid had more potent RS ratio activity than delamanid. Both in monotherapy and in combination regimens, pretomanid decreased the RS ratio more than delamanid. These results highlight that there exist significant differences between drugs that are not captured by the conventional PD marker (CFU burden). Molecular measures of *Mtb* physiology such as the RS ratio may support drug and regimen development by differentiating between the activity of drugs or regimens that have the same effect on CFU.

Conventional preclinical testing has left a state of equipoise in which it is unclear whether delamanid and pretomanid have different activity or are interchangeable.^2^ Quantifying drug activity in preclinical models has traditionally depended heavily on enumeration of *Mtb* growth on agar plates.^11^ Our previous work has suggested that measurement of properties other than pathogen burden may augment understanding of drug effects.^13,15,15–17^ Specifically, the RS ratio measures a fundamental indicator pathogen health and activity via assessment of ongoing rRNA synthesis rather than bacterial burden. We have shown that the RS ratio provides orthogonal information that is distinct from CFU ^18^ and may distinguish between drugs or regimens that have comparable bactericidal activity.^17,19^ Understanding of drug effects is often influenced by the PD marker used. Here, the traditional marker of drug effect (CFU) suggested that delamanid is significantly more potent than pretomanid whereas the RS ratio suggested the opposite.

TB drug development has been impeded by a “portability” problem in which conventional *in vitro* microbiologic measures of drug activity (*i*.*e*., minimum inhibitory concentration (MIC) and minimum bactericidal activity (MBC) generally do not translate directly to *in vivo* drug activity. Translation from *in vitro* microbiologic assays to *in vivo* results has required combinations of specialized conditions. Examples include testing in *ex vivo* caseum^21^ or conditions meant to mimic other in vivo conditions followed by integrative modeling.^22^ By contrast, the *in vitro* RS ratio activity of delamanid and pretomanid was directly concordant with their *in vivo* RS ratio activity in mice. Specifically, *in vitro* studies showed that delamanid required a higher concentration than pretomanid to suppress the RS ratio. Murine studies showed that the delamanid RS-EC_90_ was not achieved and, correspondingly, there was minimal RS ratio effect. If confirmed with additional drug classes, portability of RS ratio results from *in vitro* to mouse could de-risk and accelerate the progression from early drug discovery to animal studies. It would provide drug developers with greater confidence in advancing animal testing and enable dose-projection needed to achieve RS ratio activity.

This work has several limitations. Our *in vitro* concentrations-response analysis indicated that delamanid achieved the same RS ratio efficacy (RS-E_max_) as pretomanid if provided at sufficiently high concentration. With the intention of exceeding the *in vitro* delamanid RS-EC_90_, we treated mice with delamanid 100 mg/kg, a dose >16-times higher than human equivalent dose. However, even delamanid administered to mice at 100 mg/kg failed to reach the RS-EC_90_ concentration in plasma and, correspondingly, we observed no RS ratio effect in mice. This suggests an even higher delamanid dose would be required to achieve an RS ratio effect and highlights the relatively lower RS ratio potency of delamanid. Second, this work was performed in a single mouse model (BALB/c). Future assessment in other models, including the C3HeB/FeJ mouse, would provide added value in terms of the impact of advanced disease severity and heterogeneous pathology.^23^

This work provides a new perspective on the activity of two nitroimidazoles which play important roles in contemporary TB treatment regimens. CFU, the conventional culture-based measure of pathogen burden, showed delamanid and pretomanid had similar bactericidal activity when administered at their respective human-equivalent dose, but the RS ratio revealed that they differed in their capacity to inhibit bacterial rRNA synthesis, a fundamental physiologic process. RS ratio activity has previously been associated with treatment-shortening activity in the BALB/c mouse model,^17^ suggesting this previously unappreciated difference between delamanid and pretomanid is consequential. Given the high cost and long timelines of TB regimen development, we believe that there is risk in continuing to “put all eggs in a single [PD] basket” by focusing exclusively on CFU burden. Instead, we propose that preclinical TB drug and regimen evaluation should include not only CFU burden but also molecular measures of pathogen physiologic health.

